# Ecological Dynamics Imposes Fundamental Challenges in Microbial Source Tracking

**DOI:** 10.1101/2022.05.21.492809

**Authors:** Xu-Wen Wang, Lu Wu, Lei Dai, Xiaole Yin, Tong Zhang, Scott T. Weiss, Yang-Yu Liu

## Abstract

Quantifying the contributions of possible environmental sources (“sources”) to a specific microbial community (“sink”) is a classical problem in microbiology known as microbial source tracking (MST). Solving the MST problem will not only help us understand how microbial communities were formed, but also have far-reaching applications in pollution control, public health, and forensics. Numerous computational methods, referred to as MST solvers hereafter, have been developed in the past and applied to various real datasets to demonstrate their utility across different contexts. Yet, those MST solvers do not consider microbial interactions and priority effects in microbial communities. Here, we revisit the performance of several representative MST solvers. We show compelling evidence that solving the MST problem using existing MST solvers is impractical when ecological dynamics plays a role in community assembly. In particular, we clearly demonstrate that the presence of either microbial interactions or priority effects will render the MST problem mathematically unsolvable for any MST solver. We further analyze data from fecal microbiota transplantation studies, finding that the state-of-the-art MST solvers fail to identify donors for most of the recipients. Finally, we perform community coalescence experiments to demonstrate that the state-of-the-art MST solvers fail to identify the sources for most of the sinks. Our findings suggest that ecological dynamics imposes fundamental challenges in solving the MST problem using computational approaches.

## INTRODUCTION

Estimating the contributions or mixing proportions of different source microbial communities (“sources”) to a specific microbial community (“sink”) is known as the microbial source tracking (MST) problem^1–3^. Historically, MST was framed in the context of quantifying the input of various sources of fecal contamination to manage and remediate water pollution^4^. Recently, MST has been used in many other contexts such as healthcare^5,6^ and forensics^7^. This is largely due to the advances in metagenomics and next-generation sequencing technologies, which have enabled us to collect microbiome data at an unprecedented speed^8–11^ and provide deep insights into the roles of microbes in the integrity of their environments or the well-being of their hosts^12–14^. Despite these advances, much remains unclear regarding how the microbial communities were formed in the first place and how microbes migrate across different habitats. Understanding the origins of microbial communities by solving the MST problem is crucial for us to reveal their assembly rules, prevent future instances of contamination, and inform disease prevention.

Mathematically, the MST problem can be formalized as follows. Consider a sink community represented by a composition vector ***x***, where *x_j_* corresponds to the relative abundance of species-*j*, 1 ≤ *j* ≤ *N*. Let *K* be the number of known sources to this sink community. Each known source is represented by a composition vector ***y***^(a)^, where 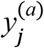 is the relative abundance of species-*j* in source-*a* (1 ≤ *a* ≤ *K*). In addition to the *K* known sources, we assume there is an unobserved source labeled as (*K* + 1). Our goal is to estimate the contributions or mixing proportions of the (*K* + 1) source communities to form the sink community, i.e., inferring *m_a_* (*a* = 1,…, *K* + 1) that satisfy 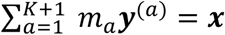 and 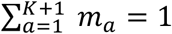.

Previous MST studies typically aimed at defining source-specific indicators (microbial or chemical) with appropriate detection techniques^1,3^. Recently, numerous computational methods based on machine learning or Bayesian modeling, referred to hereafter as MST solvers, have been developed to infer the contributions of different sources to a sink community^2,4^. Here we introduce three representative MST solvers. The first solver is based on the classification analysis in machine learning, e.g., using the Random Forest (RF) classifer^15^. In this case, each source represents a distinct class and RF will classify the sink into different classes with different probabilities. The probabilities of the sink belonging to the different classes can be naturally interpreted as the mixing proportions or contributions of those sources to the sink. Beyond the simple classification analysis, more advanced statistical methods based on Bayesian modeling have been developed. For example, SourceTracker is a Bayesian MST solver that explicitly models the sink as a convex mixture of sources and infers the mixing proportions via Gibbs sampling^16^. Due to its computational complexity, SourceTracker is only applicable to small- or medium-size datasets with a small number of sources. FEAST (fast expectation-maximization for microbial source tracking^17^) is a more recent statistical method. FEAST also assumes each sink is a convex combination of sources. But it infers the model parameters via fast expectation-maximization, which is much more scalable than Markov Chain Monte Carlo used by SourceTracker.

Both SourceTracker and FEAST have shown promising performance in synthetic datasets and offered biologically meaningful interpretations when applied to real datasets under certain contexts. Yet, the synthetic datasets used to validate these MST solvers were all generated from statistical distributions, rather than dynamics models in community ecology. Hence, the ecological dynamics driving the community assembly is completely ignored. We hypothesize that, after considering the ecological dynamics, the power of those MST solvers might be significantly restricted.

Here we consider two factors that heavily affect the ecological dynamics and community assembly: (1) microbial interactions; (2) priority effects. Microbial interactions are ubiquitous. They can be mediated by direct secretion of substances such as bacteriocins^18,19^, ecological competition between the microbes^20^, metabolite exchange^21^, or the host’s immune system modulation^22–24^. In the presence of microbial interactions, the final composition of the sink community will in general be fundamentally different from its initial one, i.e., the one right after the source mixing, which is typically not available to us (see Fig.1). Consequently, the source contributions (or mixing proportions) estimated by applying MST solvers to the final sink community will be significantly different from the source contributions estimated by applying MST solvers to the initial sink community.

**Figure 1:**
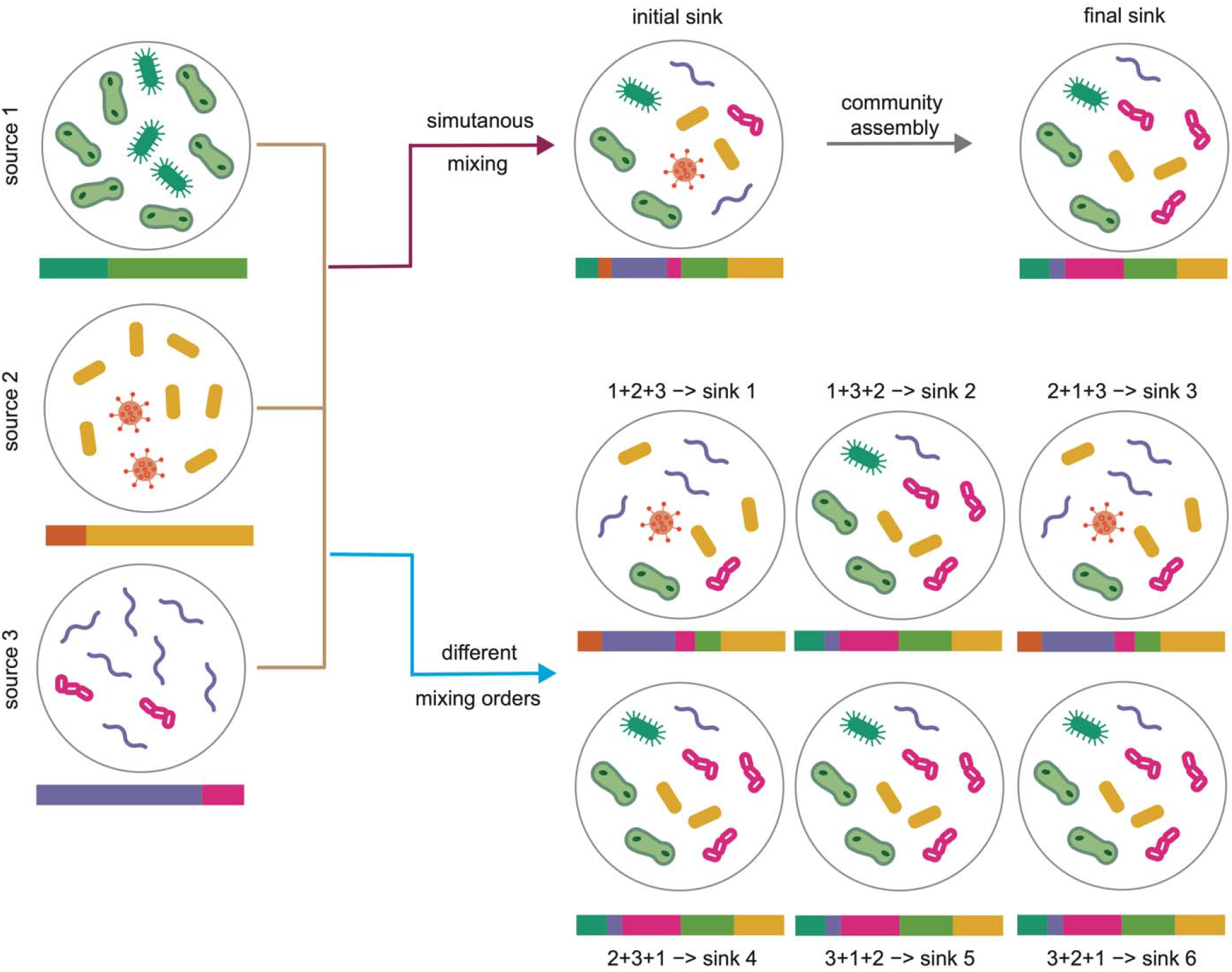
Ecological dynamics imposes fundamental challenges in microbial source tracking. (**Top**) A sink is obtained by simultaneously mixing three sources (without any species overlap) with mixing proportions (1/3,1/3,1/3). Due to the presence of microbial interactions, the initial composition of the sink community (right after the mixing, which is typically not available for MST) can be significantly different from the final composition (which is the input of MST solvers). Applying any MST solver to the final sink composition will yield different results from applying the MST solver to the initial sink composition. (**Bottom**) Due to the priority effects, three sources mixed with different orders can result in total 3! = 6 different sinks with different compositions, even if the mixing proportions of the sources are exactly the same for the different mixing orders.

Ecological theory suggests that the establishment of new species in a community can depend on the order and/or timing of their arrival, a phenomenon known as *priority effects^25–28^*. This phenomenon is actually ubiquitous in animal^29,30^, plant^31^, and microbial communities^28,32,33^. Mechanisms of priority effects and evidence for their importance have been heavily studied for microbial communities inhabiting a range of environments, including the mammalian gut^34–37^, the plant phyllosphere^38–40^ and rhizosphere^41,42^, soil^43^, freshwaters^44^ and oceans^45,46^. For example, it has been pointed out that priority effects probably shape the human gut microbiome during early childhood^47^. In particular, the infant’s exposure history and the patterns of dispersal from various sites in or on their mother could mediate the observed mutual exclusion between *Bacteroides spp., Escherichia spp*. and lactic acid producers such as *Bifidobacterium spp*. and *Lactobacillus spp*^47^. In the presence of priority effects, even if the mixing proportions (source contributions) are exactly the same, sink communities resulting from mixing the same set of sources but with different mixing orders could be drastically different (see Fig.1). Thus, for the different sink communities, the source contributions estimated by MST solvers will also be quite different, contradicting the truth.

To test our hypothesis, in this work we first systematically examined the impact of microbial interactions and priority effects on the performance of existing MST solvers using synthetic data generated by a classical population dynamics model in community ecology. We found that those solvers fail in the presence of microbial interactions or priority effects. We offered mathematical explanations for the failures. We then applied FEAST and SourceTracker, the two state-of-the-art MST solvers, to analyze data from two fecal microbiota transplantation (FMT) studies, finding that it fails to identify donors for most of the recipients. To experimentally validate our hypothesis, we performed community coalescence experiments, where fecal samples from 24 healthy individuals (i.e., sources) were mixed and cultured *ex vivo* to form 481 sink communities. We found that FEAST and SourceTracker fail to identify sources for most of the sinks. These results underscore the fundamental challenges imposed by ecological dynamics in solving the MST problem using computational approaches.

## RESULTS

### Impact of microbial interactions on MST

To illustrate the impact of microbial interactions on MST, we simulated source and sink communities as the steady states of a classical population dynamics model in community ecology --- the Generalized Lotka-Volterra (GLV) model: 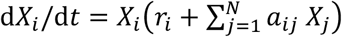, *i* = 1,…, *N*. Here *X_i_* is the abundance (or biomass) of species-*i* and *r_i_* is its intrinsic growth rate. The microbial interaction matrix 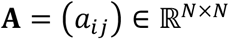 can be represented by an ecological network 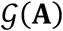: there is a directed edge (*j* → *i*) in the network if and only if *a_ij_* ≠ 0. And *a_ij_* >0 (< 0, or = 0) means that species-*j* promotes (inhibits or does not affect) the growth of species-*i*, respectively. To generate the matrix **A**, we first generate the underlying network 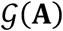 using a random graph model^48^ with *N* nodes (species) and connectivity *C* (representing the probability of randomly connecting two nodes). Then for each link 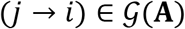 with *j* ≠ *i*, we draw *a_ij_* from a normal distribution 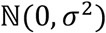. Here, the standard deviation *σ* of this normal distribution can be considered as the characteristic inter-species interaction strength. Despite its simplicity, the GLV model has been successfully applied to describe the population dynamics of various microbial communities, from the soil^49^ and lakes^50^ to the human gut^51,52^.

We generated three source communities, *S*_1_ *S*_2_ and *S*_3_, each with 30 species drawn from a pool of *N = 90* species. To simplify the MST problem, we ensured the three sources do not share any common species, and the intrinsic growth rates of all species were set to be identical (*r_i_* = 0.5 for *i* = 1,…,N). The composition vectors of *S*_1_, *S*_2_ and *S*_3_ (denoted as ***y***^(1)^, ***y***^(2)^, ***y***^(3)^, respectively) were obtained by running the GLV model until a steady state was reached and then normalizing the steady-state abundance of each species by the total biomass of the community (see SI Sec. 1 for details).

To systematically examine the impact of microbial interactions on MST, we tuned the connectivity *C* of the ecological network 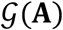 and the characteristic inter-species interaction strength *σ* in the GLV model. For a given pair of (*C*, *σ*), we simulated 100 sink communities with the initial composition vector ***x***(0) given by a random mixture of the three source communities, i.e., ***x***(0) = *m*_1_***y***^(1)^ + *m*_2_***y***^(2)^ + *m_3_****y***^(3)^, where *m_a_*’s were drawn from uniform distribution 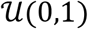 with the constraint that *∑_a_ *m*_a_* = 1. The final composition of each sink was obtained by running the GLV model until a steady state. Note that to disentangle the impacts of microbial interactions and priority effects on MST, here we assume a simultaneous mixing, i.e., all the sources (and their species) are available at the same time to avoid priority effects.

We found that, with identical intrinsic species growth rates, both FEAST and SourceTracker can achieve very high accuracy (with the coefficients of determination of the estimated proportions *R*^2^ = 1) in the absence of microbial interactions: *C* = 0 (Fig.2a) or *σ* = 0 (Fig.2b). This can be explained as follows. First, in the absence of microbial interactions and with identical intrinsic species growth rates, the final composition of each sink will be identical to its initial composition (right after the mixture of the three sources). Second, the three sources do not share any common species, hence the MST problem becomes trivial for those solvers that assume each sink is a convex combination of sources. Note that even in this ideal case, the classification-based MST solver (i.e., RF) does not perform very well. This is because, as the combination of different sources, the sink community’s composition does not necessarily need to be similar to the composition of any source.

**Figure 2:**
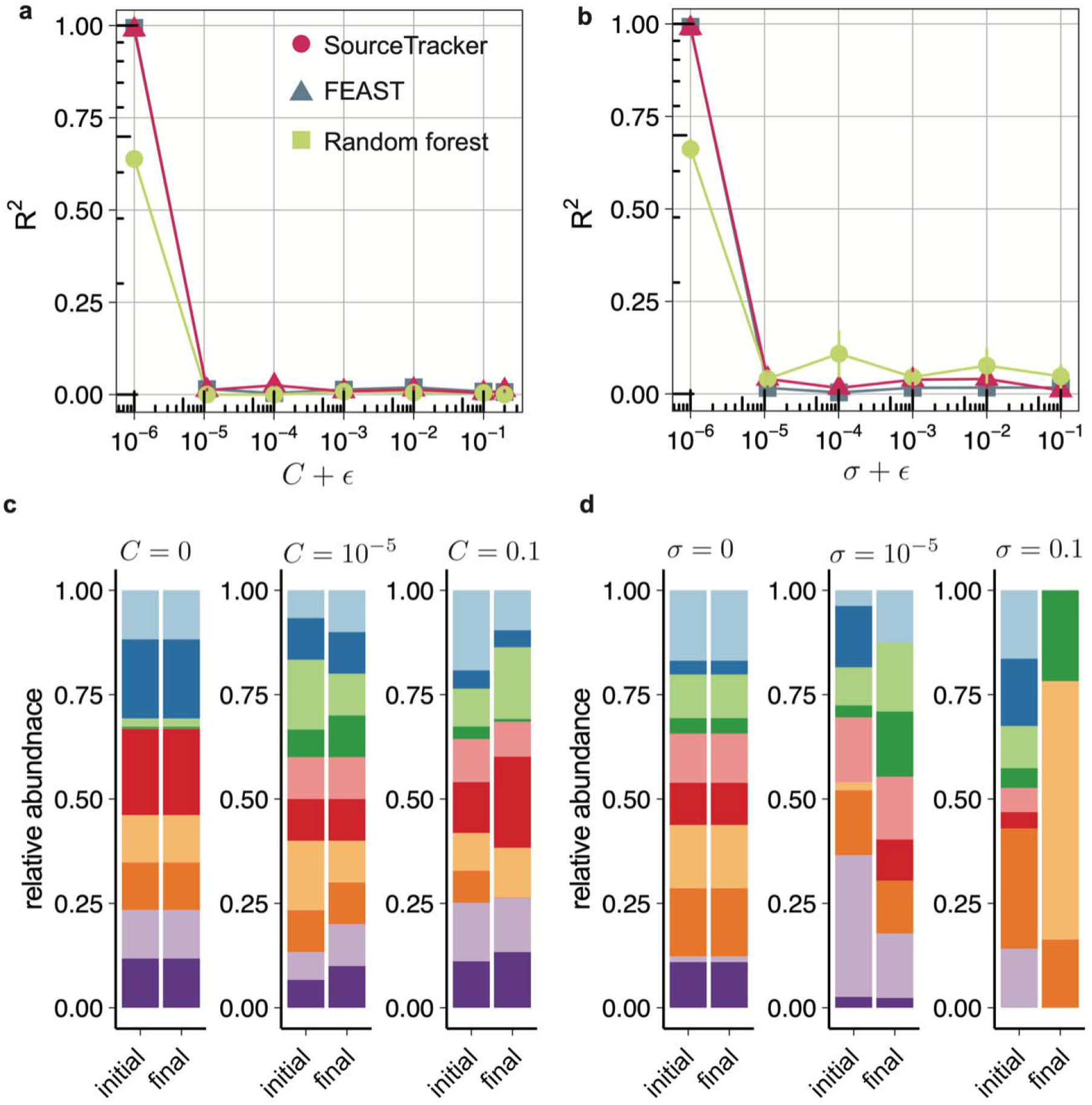
Impact of microbial interactions on MST. **a-b**, Performance of SourceTracker (red), FEAST (blue) and Random Forest (green) in simulated sinks with different network connectivity *C* (**a**) and characteristic interaction strengths *σ* (**b**). Each simulation was performed using 3 synthetic sources and 100 synthetic sinks. Accuracy of each method is measured as the coefficients of determination (*R^2^*) of the estimated proportions. Each point represents the mean *R^2^* for three independent source sets; error bars show s.e.m (*n =* 3) of the mean of *R^2^* over three sources. **c-d**, Initial and final steady compositions (we only show the relative abundance of the first 10 species for visualization purpose) of a sink with different network connectivity (**c**) and characteristic interaction strengths (**d**). In (**a,c**), the diagonal elements of the interaction matrix A are set to be *a_ii_* = −5*C* to ensure the stability of the community, and the characteristic interaction strength *σ* = 0.1. In (**b,d**), we set *a_ii_* = −5*σ* to ensure the stability, and the network connectivity *C* = 0.5. In all the simulations, we set the intrinsic growth rate *r* = 0.5 for all the species. We added a pseudo number *ϵ* = 10^−6^ to the x-axis for visualization purpose.

Interestingly, with a nonzero *C* or *σ*, none of the three MST solvers can successfully estimate the source contributions (indicated by *R^2^* ≈ 0). This implies that the existing MST solvers will completely fail as long as microbial interactions are present, and even in the absence of priority effects (see Fig.2a,b).

The unsolvability of the MST problem in the presence of microbial interactions can be conceptually explained as follows. Any microbial interactions will drive the sink community to evolve from its initial state to its final state (Fig.2c,d). The final state will be generally different from the initial one. There are two exceptions. First, the initial sink community is already at its steady state and hence will not change over time. This case almost never happens, because the initial sink is obtained by mixing multiple sources. Even though the sources are at their respective steady states, simply mixing them will not lead to another steady state. The interactions among the species across different sources will affect the assembly of the sink community. Some source-specific species might even diet out due to competition. Second, the system has a periodic trajectory in the state space, and the initial and final states happen to be identical. This coincidence generally will not happen for an unspecific time interval between the initial and final states. (See SI Sec.2 for a more mathematical explanation on the difference between the initial and final states of the sink community, using generic population dynamics models.) Since the initial and final states of the sink community are different, the source contributions estimated by applying any MST solver to the final sink community will also be different from that estimated by applying the MST solver to the initial sink community. We can avoid this issue by inferring the initial state from the final state. But this is impossible if the system is globally stable, i.e., any feasible initial state will result in the same final state. Even if such global stability does not exist, inferring the initial state from the final one would typically require detailed knowledge of the ecological dynamics, which is not known *a priori*. All these factors suggest that without *a prior* knowledge on the ecological dynamics, the MST problem is mathematically unsolvable in the presence of microbial interactions.

### Impact of priority effects on MST

To examine the impact of priority effects on MST, we again simulated three source communities *S*_1_, *S*_2_ and *S*_3_ whose species collections do not have any overlap (30 species for each source). The final compositions of sources were obtained by running the GLV model until reaching a steady state and then normalizing the steady-state abundance of each species by the total biomass of the community (see SI Sec. 1 for details). For each of the 3! = 6 mixing orders, we generated a sink by mixing three sources with equal proportion 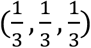, then ran the GLV model to obtain its final composition. For comparison purposes, we also generated a sink through simultaneous mixing of the three sources with equal proportion 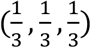. We visualized the compositions of the three sources and the seven sinks using the t-distributed stochastic neighbor embedding (t-SNE) method, finding that the compositions of the seven sinks are clearly different (see Fig.3a). We then ran FEAST, the fastest MST solver, to estimate the contributions of the three sources to each sink, finding that the contributions are different for different sinks, despite the true mixing proportions being exactly the same (Fig.3b). In the above simulations we set the network connectivity *C* = 0.5 and the characteristic interaction strength *σ* = 1.

**Figure 3:**
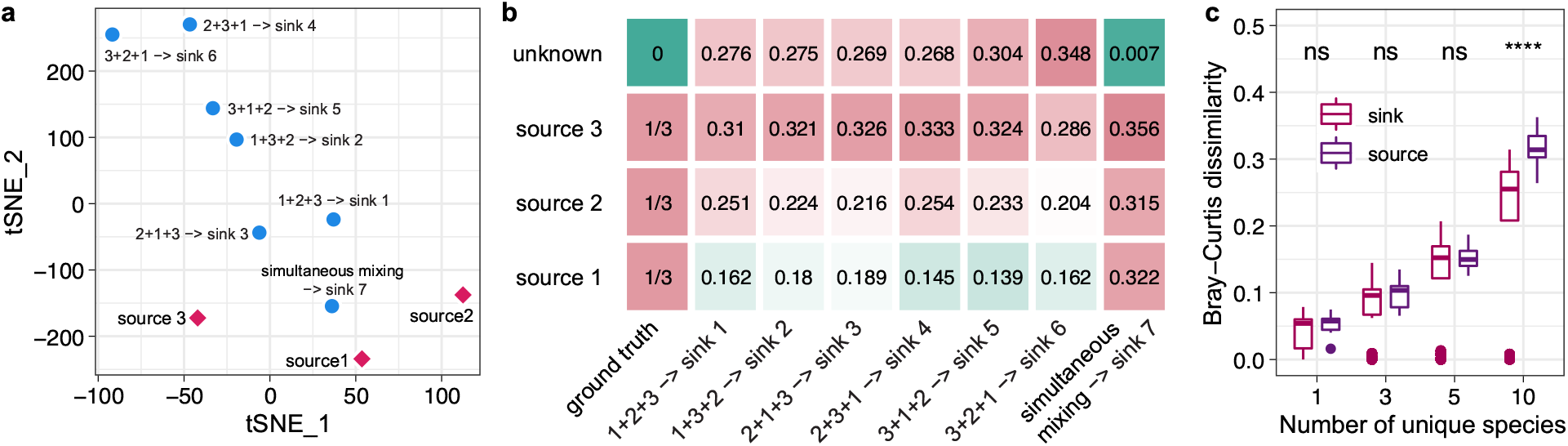
Impact of priority effects on MST. **a-b,** We synthesized three sources *S*_1_, *S*_2_ and *S*_3_ whose species collections do not have any overlap (30 species for each source). We mixed these three sources using six different mixing orders but with the same mixing proportions 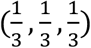, rendering six sinks. We set the network connectivity *C* = 0.5, the characteristic interaction strength *σ* = 1, the intrinsic growth rate *r* = 0.5 for each species. We set the diagonal elements of interaction matrix **A** to be *a_ii_* = −5 to ensure the stability. **a,** Dimensionality reduction using t-SNE shows the variations among the six sinks generated from the six different mixing orders. **b,** Contribution of each source to the six simulated sinks estimated by FEAST. **c,** Between-sink and between-source Bray-Curtis dissimilarity. We synthesized five sources. The species collection of each source includes *N_u_* unique species and the remaining (90 – 5A_u_) species are shared by all the sources. We mixed these five sources with the same mixing proportions 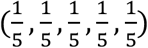 in 100 different mixing orders randomly chosen from the total 5! = 120 mixing orders. We set the network connectivity *C* = 0.5, the characteristic interaction strength *σ* = 1, the intrinsic growth rate *r* = 0.5 for each species. We set the diagonal elements of interaction matrix **A** to be *a_ii_* = −10 to ensure the stability. P-values were calculated using one-sided Wilcoxon test.

The above results make us wonder the solvability of the MST problem in the presence of priority effects. Here we offer an outline of proof that the MST problem is mathematically unsolvable in the presence of priority effects. Consider a set of source communities. If we mix them in different orders (but using the same set of mixing proportions), this will generally lead to different sink communities due to priority effects. The between-sink dissimilarity can be as large as the between-source dissimilarity (see Fig.3c). We emphasize that different mixing orders generally result in different sink communities even in the absence of any microbial interactions (see SI Sec.3 for a mathematical explanation). For different sink communities, the source contributions estimated by any computational method (i.e., MST solver) will also be different, contradicting the fact that the source contributions (i.e., mixing proportions) are exactly the same. This proof by contradiction clearly illustrates that the MST problem is mathematically unsolvable in the presence of priority effects.

### Evaluation of MST solvers using data from FMT studies

During FMT, fecal microbiota from a carefully screened, healthy donor is introduced to a recipient through either the lower or upper gastrointestinal tract. It is a “natural” mixing experiment that can be used to evaluate the performance of MST solvers. To achieve that, we applied FEAST and SourceTracker to analyze data from two FMT studies^53,54^.

In the first study, recurrent *Clostridioides difficile* infection (rCDI) patients were treated with encapsulated donor material for FMT (cap-FMT)^53^. Fig.4a shows the donor-recipient relationship between 7 healthy donors and 88 rCDI patients (i.e., recipients). Each trajectory represents a donor and one of its recipients with fecal samples collected at (up to) five different time points: pre-FMT, 2-6 days post FMT, weeks (7-20 days) post FMT, months (21–60 days) post FMT, and long term (>60 days). The Principal Coordinate Analysis (PCoA) plot of all the microbiome samples is shown in Fig.4b. We tested if FEAST can correctly identify the donor of a recipient. To achieve that, we considered each post-FMT sample of each recipient as a sink community and considered the fecal samples of all the 7 donors, as well as the recipient’s pre-FMT sample as potential source communities. Then we applied FEAST to solve the MST problem. For each sink community, among all the 7 donors, we referred to the one whose fecal sample has the highest contribution estimated by FEAST as the “predicted donor” (green squares, Fig.4c, Fig.S1). Interestingly, we found that for a large portion (61%) of the sink communities, FEAST failed to identify the true donor (red circles, Fig.4c, Fig.S1), though the average Jensen-Shannon divergence among those donors is higher enough (0.63). Similar results were found for SourceTracker (see Fig.S2). These results clearly demonstrate the limitation of existing MST solvers.

**Figure 4:**
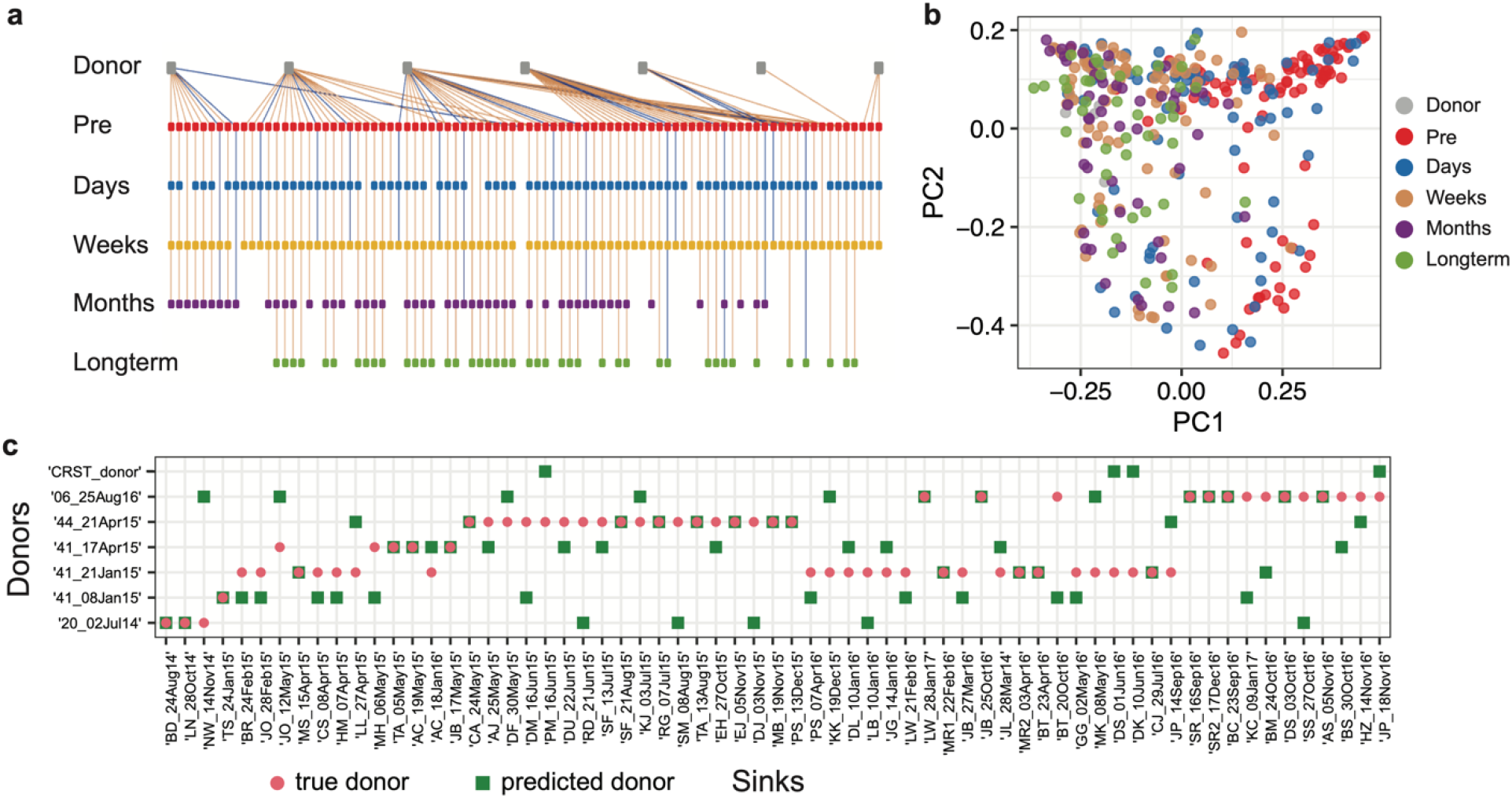
Evaluation of FEAST using FMT data from Staley et al.^53^. **a,** Donor-recipient relationship. Each trajectory represents a donor and its corresponding recipients at up to 5 time points. Trajectories of recipients who responded to FMT (i.e., responders) are colored in yellow. Trajectories of non-responders are colored in blue. **b,** Principal Coordinates Analysis (PCoA) plot based on the Bray-Curtis dissimilarity. **c,** True donor (red cycle) vs. predicted donor (green square) of each recipient. For each post-FMT community (sink), among all the 7 donors, we referred to the one whose fecal sample has the highest contribution estimated by FEAST as the “predicted donor”. Here, we only showed the results for the first 65 sinks for the visualization purpose (see Fig.S1 for results of the remaining 194 sinks). Sources: microbiome samples of donors and the pre-FMT samples of recipients; Sinks: post-FMT samples of recipients.

In the second FMT study, the gut microbiota of human donors with autism spectrum disorder (ASD) or typically-developing (TD) controls were transplanted into germ-free mice^54^. The dataset includes 8 donors, 13 recipients, and in total 106 post-FMT sink communities. We again examined whether FEAST can correctly identify the true donor of each sink community. For each sink community, among the 8 donors, we refer to the one whose fecal sample has the highest contribution predicted by FEAST as the “predicted donor” (green squares, Fig.S3). We found that for 40% of the sink communities, FEAST failed to identify the true donor (red circles, Fig.S3). Similar results were observed for SourceTracker (see Fig.S4).

### Evaluation of MST solvers using data from community coalescence experiments

To further evaluate MST solvers using real data, we performed community coalescence experiments, where fecal microbiota from 24 healthy individuals (i.e., sources) were mixed and cultured *ex vivo* to form 481 sink communities (see SI Sec.4 for details). Among the 481 sinks, 256 sinks were obtained by mixing two different sources (pair-wise mixing), and the remaining 225 sinks were obtained by mixing four different sources (quadruple-wise mixing). After inoculation, the sink communities were transferred into fresh medium every 24 hours (1:200 dilution) for 10 transfers^55^ (see Fig.5a). Samples collected at the final time point were sequenced and the resulting taxonomic profiles were considered as the steady-state composition of sinks (see Methods). As expected, we found that the source and sink communities had distinct taxonomic profiles (Fig.S5–S6).

**Figure 5:**
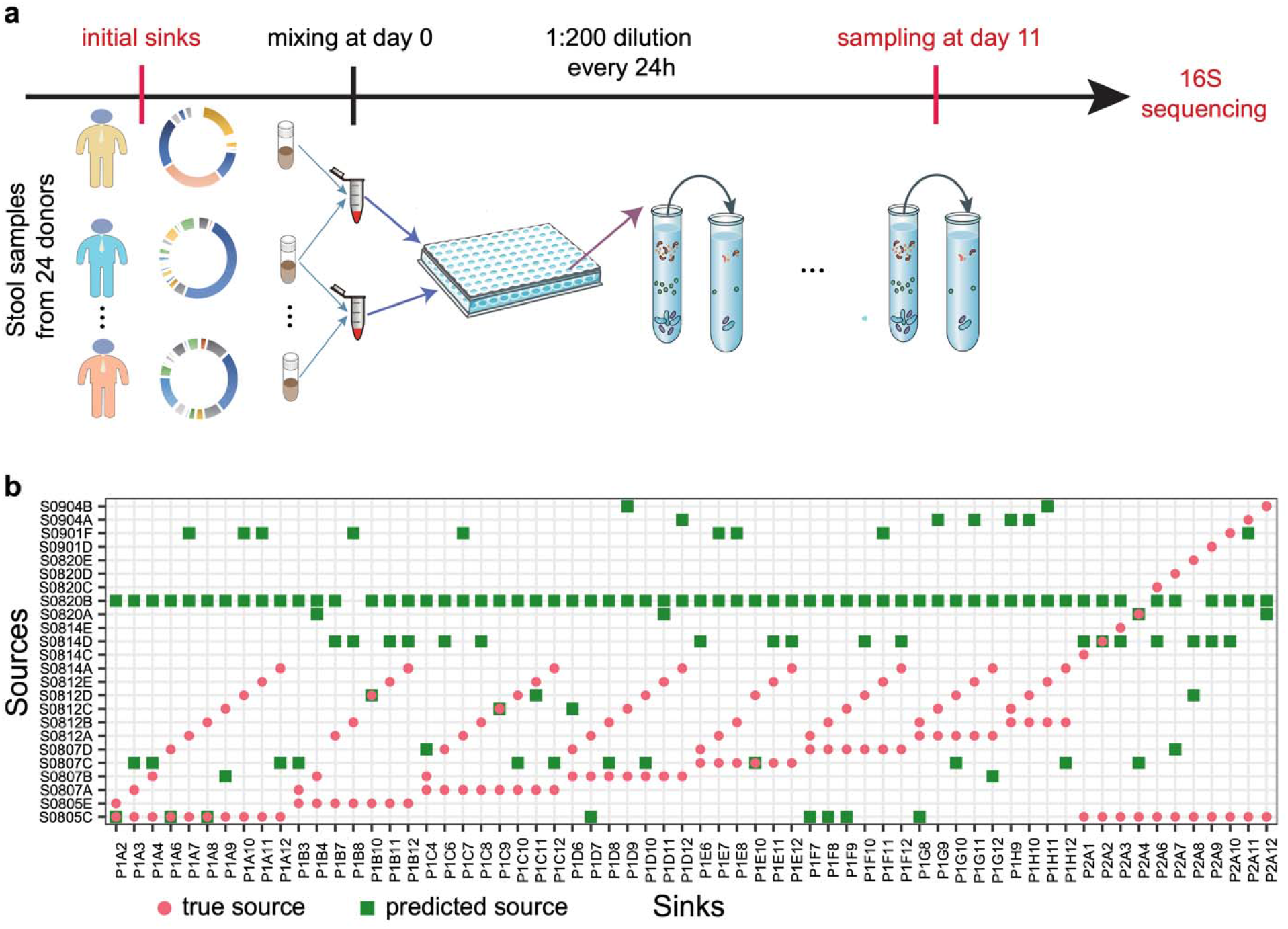
Evaluation of FEAST using data from pairwise community coalescence experiments. **a,** Schematic diagram of the community coalescence experiments. There are 24 source communities (stool samples from 24 healthy individuals). Each sink community is obtained by mixing two different source communities *ex vivo* and the final composition of each sink was obtained from metagenomic sequencing of samples collected after 11 days of the *ex vivo* mixing. **b,** True sources (red cycles) vs. predicted sources (green squares) of each sink. For each sink, among the 24 known sources, the two sources with the top-two largest contributions predicted by FEAST were referred to as the predicted sources. Here, we only showed the first 64 sinks for the visualization purpose (see Fig.S5 for results of the remaining 192 sinks).

To examine the performance of FEAST in community coalescence experiments, we first applied FEAST to analyze the compositions of the 256 sinks obtained in the pair-wise mixing experiments. We ranked the estimated contributions of 24 potential sources to each sink and selected the top-two as the predicted sources. We found that the predicted sources (green squares) are different from the true sources (red circles) for most of the 256 sinks (Fig.5b and Fig.S7). This is also true for the cases of quadruple-wise mixing (Fig.S9). Similar results were observed for SourceTracker (see Fig.S8, S10).

Note that some donor samples (e.g., S0820B, S0814D) were predicted as sources for many sinks. We found this is due to the high abundance of common ASVs shared by sinks and those particular sources (Fig.S11).

## DISCUSSION

Many computational methods have been developed to solve the MST problem. Yet, those methods ignored the underlyng ecological dynamics that drive the assembly of microbial communities. For example, as a Bayesian MST solver, SourceTracker explicitly models the sink as a convex mixture of sources and infers the mixing proportions via Gibbs sampling^16^. This approach was inspired by the “analogy” between quantifying the proportion of different source environments to a sink microbial community and inferring the mixing proportions of conversation topics in a test document^56,57^. Here we point out that this analogy is inappropriate. In topic modeling, which is a specific research area in natural language processing, the goal is to discover the abstract “topics” that occur in a collection of documents. In a sense, those documents are static or “dead”. By contrast, in MST we are typically dealing with alive (or even flourishing) microbial communities, where ecological dynamics plays an important role in community assembly and determining their state, i.e., the microbial composition. In the presence of ecological dynamics, a sink community cannot be simply considered as a convex mixture of known and unknown sources. In this work, through numerical simulations, analytical calculations, and real data analysis, we presented compelling evidence that ecological dynamics impose fundamental challenges in MST. In particular, we clearly demonstrate that the presence of either microbial interactions or priority effects will render the MST problem mathematically unsolvable for any MST solver.

MST solvers have been applied to various real datasets and demonstrated their utility across two fundamentally different contexts. First, as originally intended, they were used to quantify the contribution of different source environments to a sink microbial community. For example, SourceTracker was used to estimate the contributions of bacteria from ‘gut’, ‘oral’, ‘skin’, ‘soil’ and ‘unknown’ sources to several indoor sink environments (e.g., office buildings, hospitals, and research laboratories)^16^. It was found that wet-lab surface communities tended to be composed mainly of bacteria from ‘skin’ and ‘unknown’, while neonatal intensive care units and office communities were typically dominated by skin bacteria. FEAST was used to estimate if taxa in the infant gut originate from the birth canal, or if they are derived from some other external source at a later time point^17^. By treating samples taken from the infants at age 12 months as sinks, considering respective earlier time points and maternal samples as sources, a significantly larger maternal contribution in vaginally delivered infants over cesarean-delivered infants was found. Moreover, biological mothers were more likely to be identified as sources of their infant’s microbiome than other potential source communities. Although these results seem reasonable and agree well with our intuition, we suggest that the whole community of microbiome research should be very cautious when interpreting the results of existing MST solvers in this context. The source contributions estimated by MST solvers might be quite different from the true contributions due to complex ecological dynamics. This is particularly important for microbial communities living in nutrient-rich environments such as the human gut. For microbial communities living in oligotrophic environments (e.g., soil, ocean, etc.), the growth rates of bacteria and assembly process of communities are relatively slow^58,59,60^ and the impact of ecological dynamics on MST might be relatively low^61^. But even in this case, interpreting the results of existing MST solvers should be done with great caution.

Second, MST solvers have been used as a metric of similarity^17^. In this context, instead of quantifying the contribution of different sources to a sink, we aim for capturing the similarities between the sink and its characteristic environments using mixing proportions estimated by MST solvers. Each sink can be represented by a similarity feature vector, characterizing its similarity to each of its characteristic environments. For example, FEAST has been used in this context to distinguish patients in ICU from healthy adults, and capture shifts in microbial community composition that may underlie differences between pathogenic and neutral phenotypes^17^. We think this is a much more meaningful and practical way of using MST solvers to analyze real data.

A recent study has shown that the strain tracking approach^62^ can predict whether two metagenomics samples originate from the same donor via counting the number of species that share closely-related strains. Yet, the contribution of different sources to a given sink remains unknown. More importantly, challenges imposed by ecological dynamics are still there, which do not rely on a particular sequencing method. For example, in the presence of microbial interactions and priority effects, those source-specific microbial strains may not be able to survive in the sink community at all. This actually raises a serious concern on any approaches based on indicator species in solving the MST problem.

## Data and code availability

The sequencing data from the first FMT study is available at Sequence Read Archive at the National Center for Biotechnology Information under BioProject accession number SRP070464. The sequencing data from the second FMT study is available at Qiita^63^ with ID: 11809. The raw sequencing data from the community coalescence experiments is available at European Nucleotide Archive (ENA) under study accession number PRJEB51290. The code used to generate the simulated data is available at: https://github.com/spxuw/MST.

## Author Contributions

Y.-Y.L conceived and designed the project. X.-W.W and Y.-Y.L did the analytical calculations. X.-W.W did all the numerical calculations and analyzed all the simulated and real datasets. L.W. and L.D. designed and performed the community coalescence experiments. X.-W.W. and Y.-Y.L wrote the manuscript. L.W., L.D., X.Y., T.Z., and S.T.W interpreted the results, reviewed and edited the manuscript. All authors approved the manuscript.

## Acknowledgement

Y.-Y.L. acknowledges grants from National Institutes of Health (R01AI141529, R01HD093761, RF1AG067744, UH3OD023268, U19AI095219 and U01HL089856).

## Supplementary Information

### 1. Using an ecological model to generate synthetic microbiome data

To systematically reveal the impacts of the microbial interactions and priority effects on MST, we generated synthetic data using the classical Generalized Lotka-Volterra (GLV) model^1^:

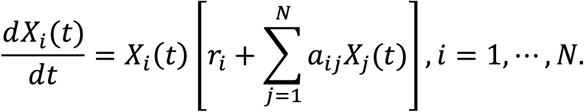

Here *X_i_*(*t*) represents the absolute abundance of species-*i* at time *t* ≥ 0, *r_i_* is its intrinsic growth rate, which is randomly drawn from a uniform distribution 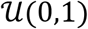, if not specified otherwise. The inter-species interactions are encoded in the interaction matrix 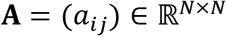, with *α_ij_* > 0 (< 0, or = 0) means that species-*j* promotes (inhibits or does not affect) the growth of species-*i*, respectively. To generate the matrix **A**, we first generate the underlying ecological network 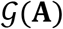 using an Erdős-Rényi random graph model^2^ with *N* nodes (species) and connectivity *C* (the probability of randomly connecting two nodes). Then for each link 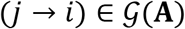 with *j* ≠ *i*, we draw *α_ij_* from a normal distribution 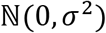. The standard deviation *σ* of this normal distribution represents the characteristic inter-species interaction strength. To ensure the stability of the system, the diagonal elements of **A** are set to be *α_ii_* = −*dC* in tuning *C* or *α_ii_* = −*dσ* in tuning *σ*. Here *d* is a positive constant. All other entries of **A** are set to be zero.

We generated *k* source communities, *S*_1_, *S*_2_,…, *S_k_* each with *N*_s_ species drawn from a pool of *N* = 90 species. To simplify the MST problem, the intrinsic growth rates of all species were set to be identical (*r* = 0.5). The composition vectors of *S*_1_, *S*_2_, *S_k_* (denoted as ***y***^(1)^, ***y***^(2)^,…, ***y***^(k)^, respectively) were obtained by running the GLV model (i.e., numerically solving the ordinary differential equations (ODEs) in the GLV model) with initial species abundances randomly chosen from a uniform distribution 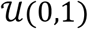, until a steady state was reached and then normalizing the steady-state abundance of each species by the total biomass of the community.

The sink obtained by simultaneously mixing the *k* sources was simulated as follows:

1. The mixing proportions of *k* sources were randomly drawn from a uniform distribution with constraint 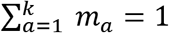.
2. The initial composition of the sink community is calculated as: ***x***(0) = *m*_1_***y***^(1)^ + *m*_2_***y***^(2)^ +…+ *m_k_**y***^(*k*)^. And the initial (absolute) abundance vector is chosen to be ***X***(0) = ***x***(0).
3. Run the GLV model until it reaches a steady state and normalize the steady-state abundance vector by the total biomass of the sink community to get the final composition of the sink community.

Consider a particular mixing order *π* among the total *k*! mixing orders. Let *π*(*α*) denote the label of the *α*-th source in the mixing order. *α,π*(*α*) ∈ {1,…, *k*}. The sink obtained by mixing the *k* sources in the order *π* was simulated as follows:

1. The mixing proportions of the *k* sources were set to be equal: 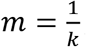,*a* = 1,…, *k*.
2. The initial abundance vector of the sink community is determined by the composition of the first source in the order *π*, i.e., *π*(1), as 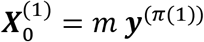. Then we run the GLV model until it reaches a steady state. Denote the steady-state abundance vector as.
3. Then the second source *π*(2) arrives. Right after the mixing, the abundance vector of the sink community becomes 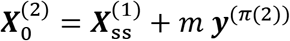. Then we run the GLV model until it reaches a steady state. Denote the steady-state abundance vector as 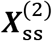.
4. Repeat step-3 until all the *k* sources have been added to the sink. Note that right after the arrival of the *k*-th source, the abundance vector of the sink community becomes 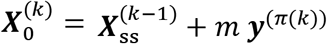. Then we run the GLV model until it reaches a steady state. Denote the steady-state abundance vector as 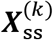.
5. Normalize the final steady-state abundance vector 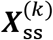 by the total biomass of the sink community to get the final composition of the sink community.

Since the input data of MST solvers is the OTU count table, for both sink and source communities, we converted the species relative abundances into counts by multiplying the absolute abundances and a fix number (1,000 in all the simulations) and rounding to the nearest integers as the synthetic count data generated by the GLV model.

### 2. Microbial interactions affect the assembly of the sink community

The deep reason why the existing MST solvers are almost doomed to fail in the presence of microbial interactions is that the true contributions of different sources are only reflected in the sink’s initial composition, which will evolve to a final composition following complex ecological dynamics. In general, the final composition will be quite different from the initial one. Here we sketch a proof.

Let us consider a sink generated by mixing *K* non-overlapping sources with compositions given by ***y***^(1)^,…, ***y***^(*K*)^, respectively. The initial abundance vector of the sink is denoted as ***X***(0) = (*X*_1_(0),…, *X_N_*(0)), and its initial composition is given by ***x***(0) = (*x*_1_(0),…, *x_N_*(0)) with 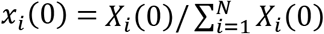 representing the relative abundance of species *i*. Note that ***x***(0) = **∑_*a*_***m_a_ **y***^(*a*)^. Let’s assume the population dynamics of the sink community can be represented by a set of ordinary differential equations:

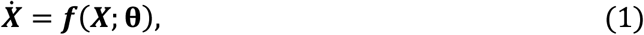

where ***X***(*t*) = (*X*_1_(*t*),…, *X_N_*(*t*)) represents the abundance vector at time *t*, ***f*** is an unspecified nonlinear function with **θ** encoding all the ecological parameters, i.e., intrinsic growth rates, and intra- and inter-species interaction strengths. After a small time-step *δt*, the abundance vector of the sink can be approximated as ***X***(*δt*) = ***X***(0) + *δt* ***f***(***X***(0); **θ**). The ratio of relative abundance for any species pair (*i, j*) in the initial community is 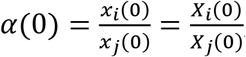, while after *δt* the ratio becomes:

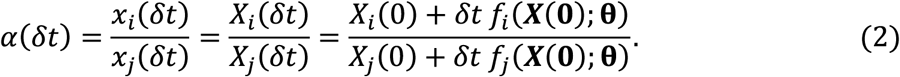

If *X_i_*(0) = *X_j_*(0) and *f_i_*(***X***(**0**); **θ**) = *f_j_*(***X***(**0**); **θ**), then we have *α*(*δt*) = *α*(0). But the condition *X_i_*(0) = *X_j_*(0) is too strong to be true. If *X_i_*(0) ≠ *X_j_*(0), but *f_i_*(***X***(**0**); **θ**) = *X_i_*(0)*g_i_*(**θ**) and *g_i_*(**θ**) = *g_j_*(**θ**), then we have *α*(*δt*) = *α*(0). For a general population dynamics model, this requirement means that there are no inter-species interactions and the intrinsic growth rates of different species are identical, which is also too strong to be true. Hence, in general *α*(*δt*) ≠ *α*(0), and the final composition of the sink community will be quite different from its initial composition.

### 3. Priority effects affect the assembly of the sink community

Consider three source communities *S*_1_, *S*_2_ and *S*_3_. Let’s assume species-*i* is only present in the source and species-*j* is only present in the source *S_2_*. We mix the source communities in 6 different orders but with identical mixing proportions 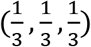. For each mixing order, we assume the arrival time of the three sources are 0, *τ*, 2*τ*, respectively, where *τ* is a constant (and is large enough for the resulting sink community to reach a steady state). Suppose we use the composition of the final sink community taken at 3*τ* to estimate the contribution of each source. We want to prove that at time *t* = 3*τ*, the sink communities resulting from different mixing orders will have different compositions, even in the absence of any microbial interactions. To achieve that, let’s compute the ratio between the relative abundance of species-*i* and that of species-*j* in the final sink community at time *t* = 3*τ*, i.e., 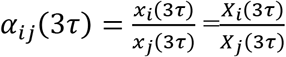.

Consider a particular mixing order *S*_1_ → *S*_2_ → *S*_3_. In the absence of any inter- or intra-species interactions, species will grow exponentially. Hence, at time *t* = 3*τ*, the abundance of a species-*i* (which is only present in the source *S*_1_) is given by: *X_i_*(3*τ*) = *m*_1_*X_i_*(0) exp(3*τr_i_*), where 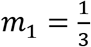 is the mixing proportion (contribution) of the source *S*_1_, *X_i_*(0) is the initial abundance of species-*i* in the source *S*_1_, *r_i_* is the intrinsic growth rate of species-*i*. Similarly, at time *t* = 3*τ*, the abundance of species-*j* (which is assumed to be only present in the source *S*_2_) is given by: *X_j_*(3*τ*) = *m*_2_*X_j_*(0)exp(2*τr_j_*). So, we have

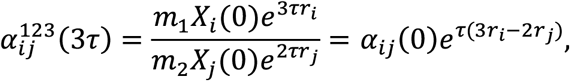

where the superscript ‘123’ indicates the mixing order *S*_1_ → *S*_2_ → *S*_3_. We can repeat the above calculation for different mixing orders. The results are summarized here:

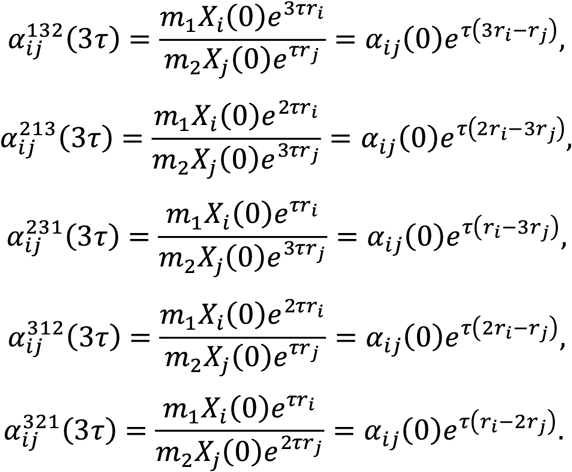

Note that if the three sources were mixed simultaneously, then we have

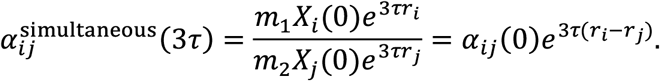

Therefore, even in the absence of any microbial interactions, different mixing patterns will result in different final compositions of the sink community, which are also different from that obtained by simultaneous mixing.

### 4. Community coalescence experiments

Stool samples from healthy human donors were collected and immediately transferred into the anaerobic workstation (85% N_2_, 10% H_2_ and 5% CO_2_, COY). 10g stool samples were suspended into 50 mL 20% glycerol (in sterile phosphate-buffered saline, with 0.1% L-cysteine hydrochloride). The samples were homogenized by vortexing and then filtered with sterile nylon mesh to remove large particles in fecal matter. Aliquots of the suspension were placed in sterile cryogenic vials and frozen at −80 °C for long-term storage until use.

Stool samples of 24 individuals were used for the community coalescence experiments. To generate 481 sink communities, samples from two, three or four different individuals were mixed with equal volume. 20 uL stool mixture was inoculated into 980 uL medium in 96-well plates (PCR-96-SG-C, Axygen) for static culturing at 37 °C in the anaerobic workstation. The medium used for *ex vivo* culture was modified from previous studies, which comprises: peptone water (2.0 g/L, CM0009, Thermo Fisher), yeast extract (2.0 g/L, LP0021B, Thermo Fisher), L-cysteine hydrochloride (1 g/ L), Tween 80 (2 mL/L), hemin (5 mg/L), vitamin K1(10 μL/L), NaCl (1.0 g /L), K_2_HPO_4_ (0.4 g/L), KH_2_PO_4_ (0.4 g/L), MgSO_4_·7H_2_O (0.1 g/L), CaCl_2_·2H_2_O (0.1 g/L), NaHCO_3_ (4 g/L), porcine gastric mucin (4 g/L, M2378, Sigma-Aldrich), sodium cholate (0.25 g/L) and sodium chenodeoxycholate (0.25 g/L)^3^. Ex vivo culture of gut microbial communities was transferred into fresh medium every 24h (1:200 dilution), for a total of 10 transfers. After each transfer, samples were centrifuged to remove the supernatant and the pellets were stored at −80°C with a plastic seal until DNA extraction.

The initial stool and *ex vivo*-cultured samples after 10 passages were sequenced. For stool samples, DNA was extracted using the QIAamp Power Fecal Pro DNA Kit (Qiagen) according to the manufacturer’s instructions. For cultured samples, DNA extraction (DNeasy UltraClean 96 Microbial Kit, Qiagen) and 16S amplicon library preparation were performed by an automated protocol at Tecan Freedom EVO 200. V3-V4 region of 16S rRNA gene was amplified using primers 341F 5’-CCTACGGGNGGCWGCAG −3’ and 805R 5’-GACTACHVGGGTATCTAATCC-3’ with custom barcodes^4^. Libraries were further pooled together at equal molar ratios and sequenced by Illumina NovaSeq (250 bp paired-end reads) at Novogene Technology (Tianjin, China).

16S amplicon sequencing data were analyzed by QIIME2 (version 2020.2)^5^. Primers of the raw sequence data were cut with Cutadapt (via q2-cutadapt)^6^. Quality control was performed by DADA2 (via q2-dada2)^7^. All amplicon sequence variants (ASVs) from DADA2 were used to construct a phylogenic tree with fasttree2 (via q2-phylogeny)^8^. The ASVs were assigned to taxonomy with naïve Bayes classifier (via q2-feature-classifier)^9^ against the SILVA database (SILVA_132_SSURef_Nr99). The ASV table was normalized, and rare ASVs (all features with a total abundance of less than 10 and present in only a single sample) were filtered out.

---

**Figure S1:**
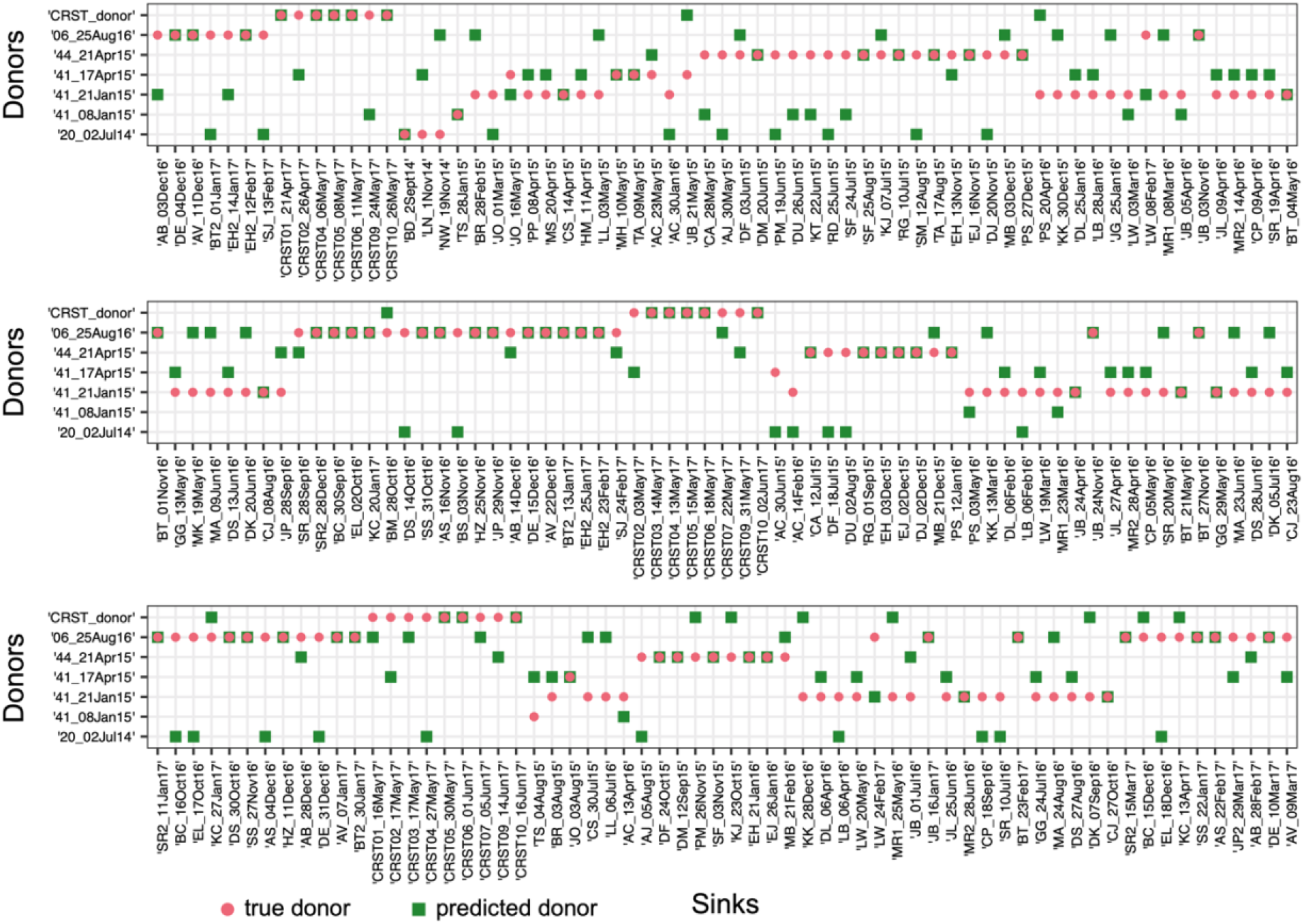
Evaluation of FEAST using FMT data from Staley et al.^10^. True donor (red cycle) vs. predicted donor (green square) of each recipient. For each post-FMT community (sink), among all the 7 donors, we referred to the one whose fecal sample has the highest contribution estimated by FEAST as the “predicted donor”. In Fig.4c, we presented results of the first 65 sinks. Here, we showed the results of the remaining 194 sinks. Sources: microbiome samples of donors and the pre-FMT samples of recipients; Sinks: post-FMT samples of recipients.

**Figure S2:**
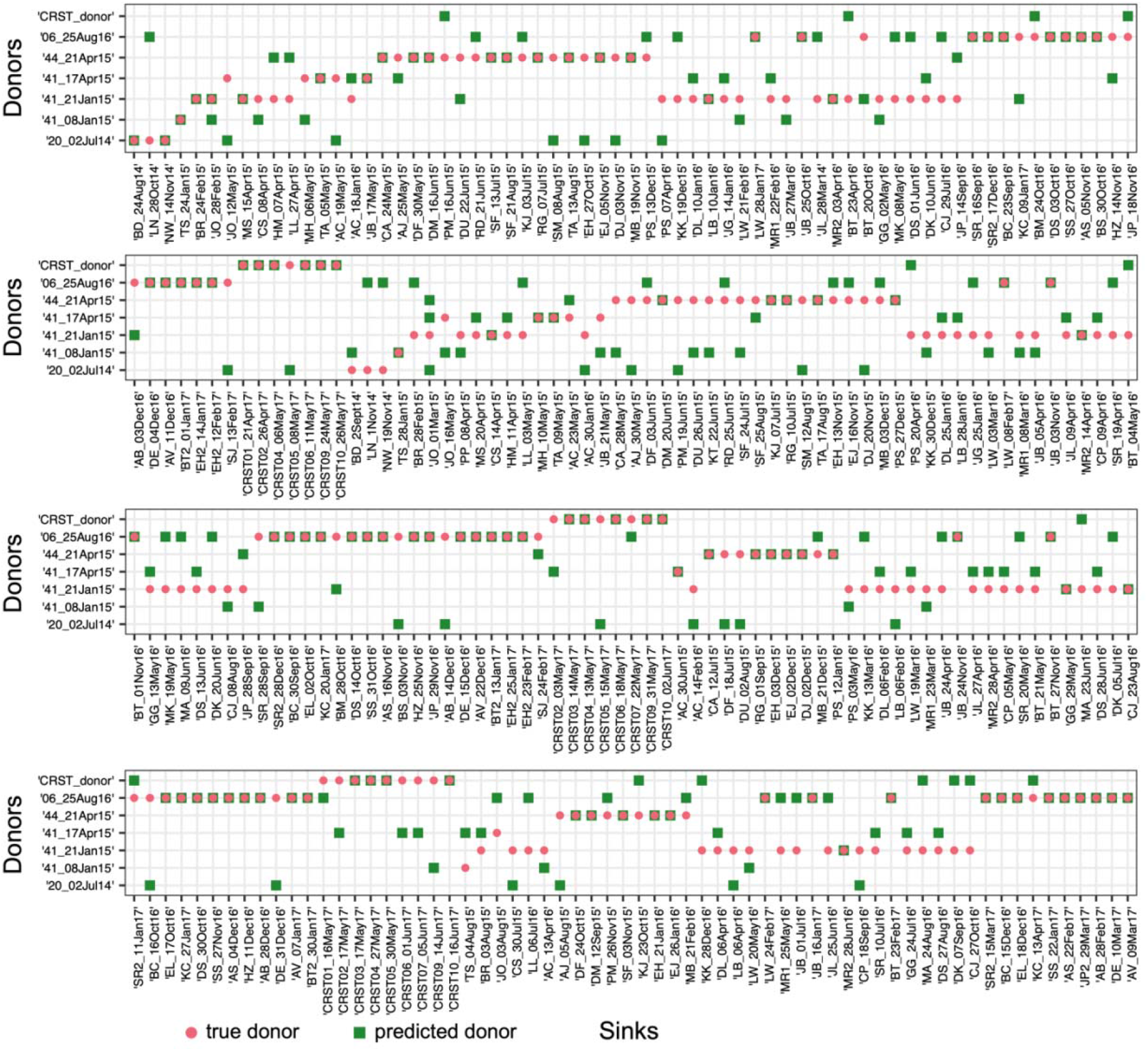
Evaluation of SourceTracker using FMT data from Staley et al.^10^. True donor (red cycle) vs. predicted donor (green square) of each recipient. For each post-FMT community (sink), among all the 7 donors, we referred to the one whose fecal sample has the highest contribution estimated by SourceTracker as the “predicted donor”. Sources: microbiome samples of donors and the pre-FMT samples of recipients; Sinks: post-FMT samples of recipients.

**Figure S3:**
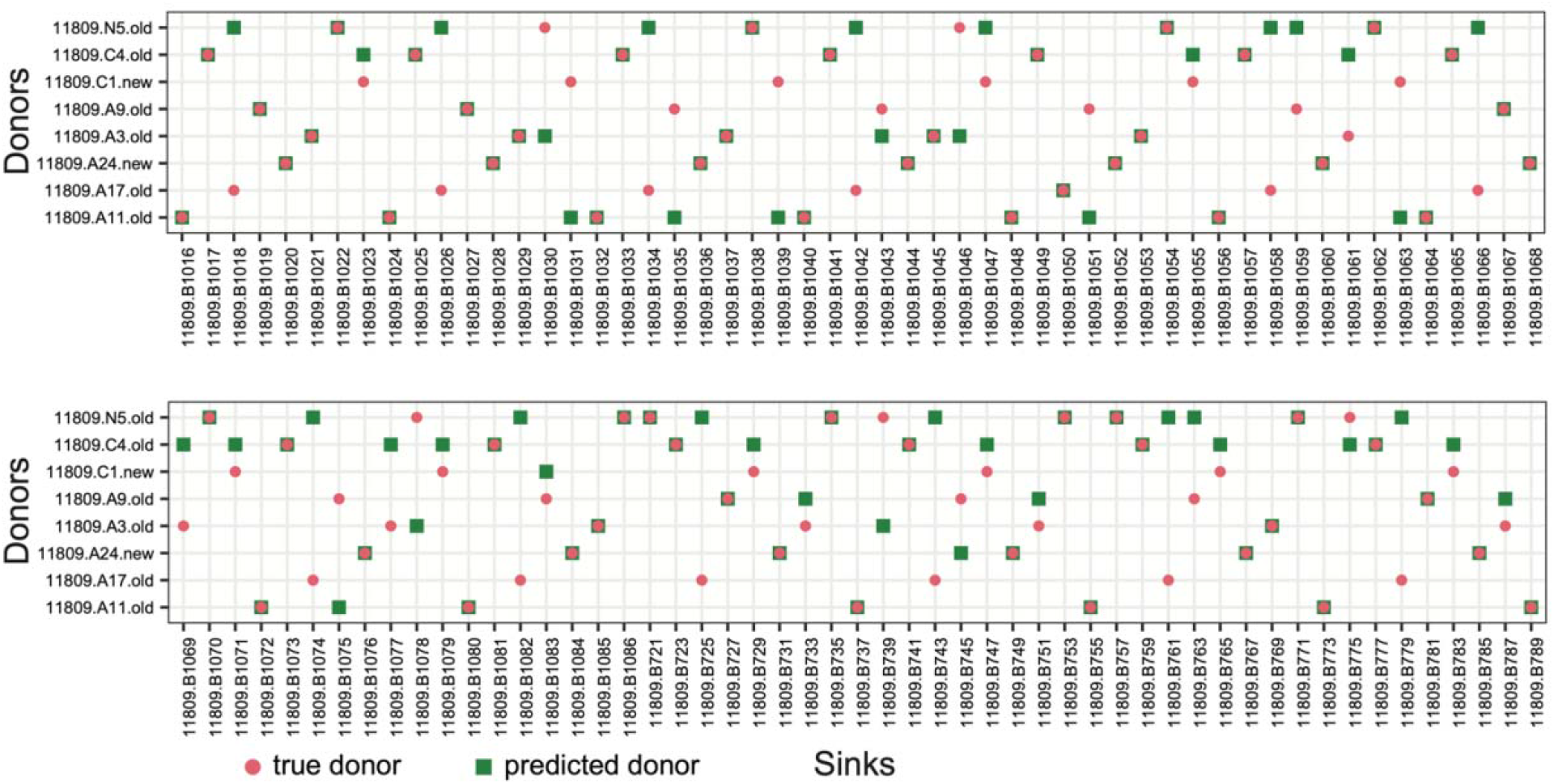
Evaluation of FEAST using FMT data from Sharon et al.^11^. True donors (red cycle) vs. the predicted donor (green square) of each recipient sink given by FEAST using the source and sink compositions as the input. For each post-FMT community (sink), among all the 8 donors, we referred to the one whose fecal sample has the highest contribution estimated by FEAST as the “predicted donor”. Sources: microbiome samples of donors and the pre-FMT samples of recipients; Sinks: post-FMT samples of recipients. In total, there are 106 sinks.

**Figure S4:**
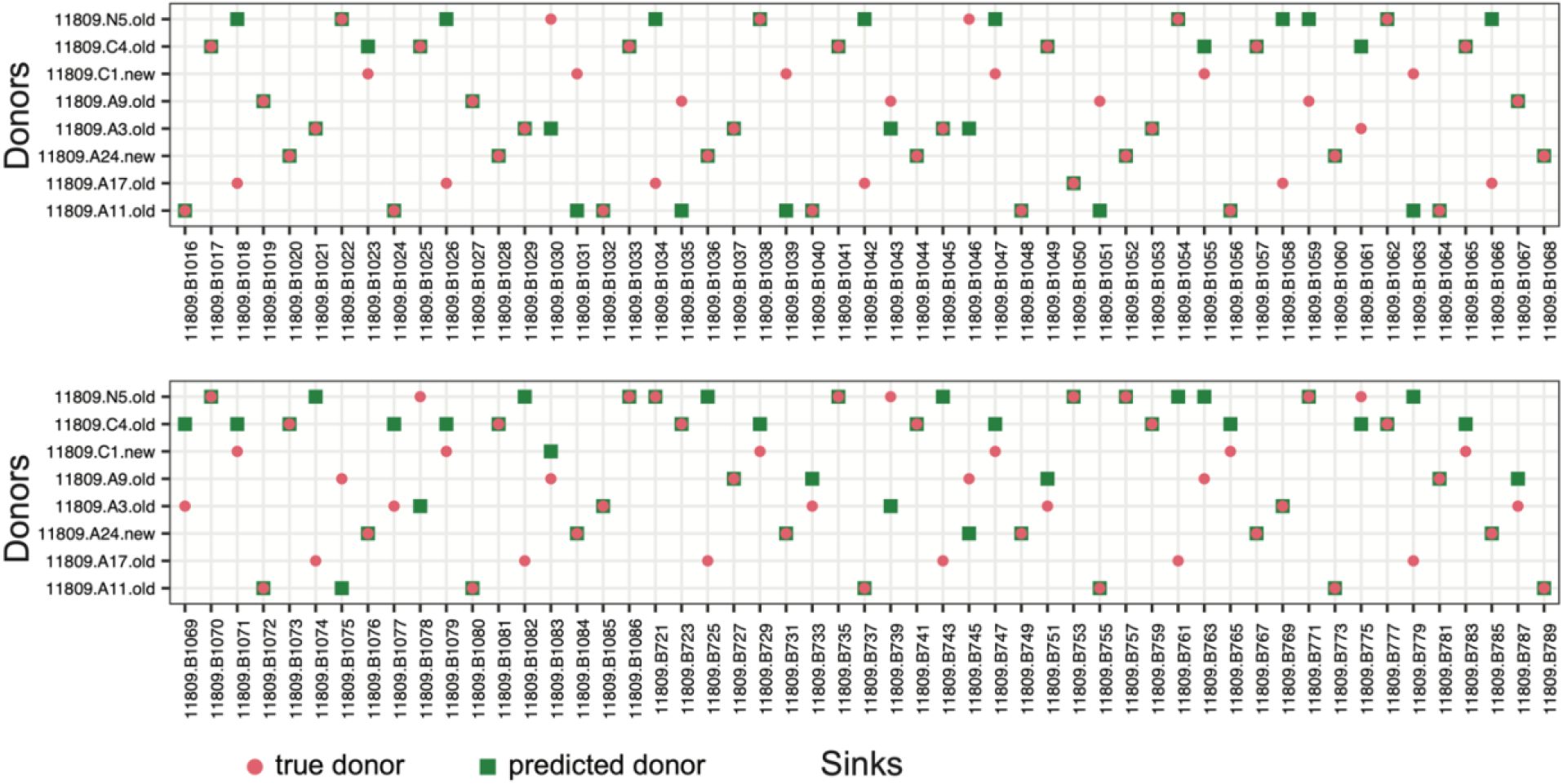
Evaluation of SourceTracker using FMT data from Sharon et al.^11^. True donors (red cycle) vs. the predicted donor (green square) of each recipient sink given by SourceTracker using the source and sink compositions as the input. For each post-FMT community (sink), among all the 8 donors, we referred to the one whose fecal sample has the highest contribution estimated by SourceTracker as the “predicted donor”. Sources: microbiome samples of donors and the pre-FMT samples of recipients; Sinks: post-FMT samples of recipients. In total, there are 106 sinks.

**Figure S5:**
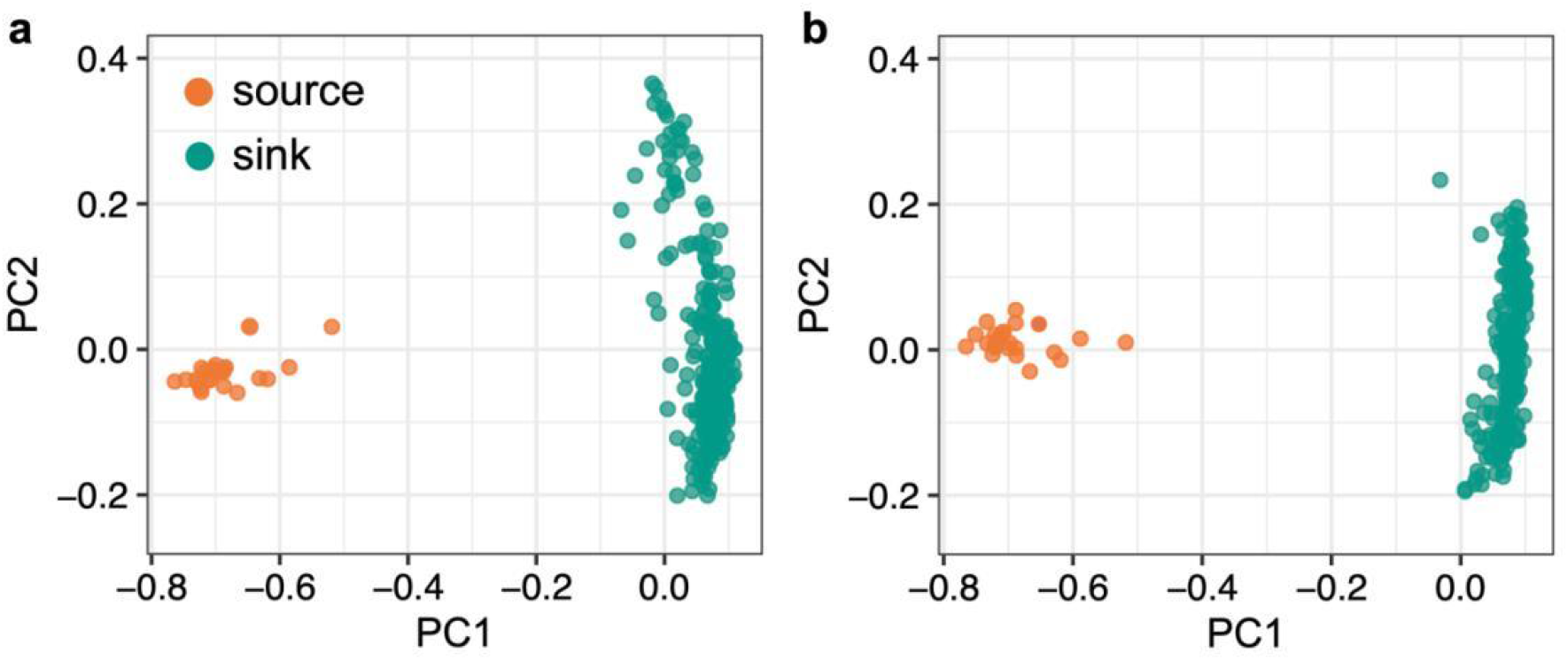
Principal Coordinate Analysis (PCoA) plot of the sinks and sources in the community coalescence experiments. **a**, Pairwise mixing. **b**, Quadruple-wise mixing.

**Figure S6:**
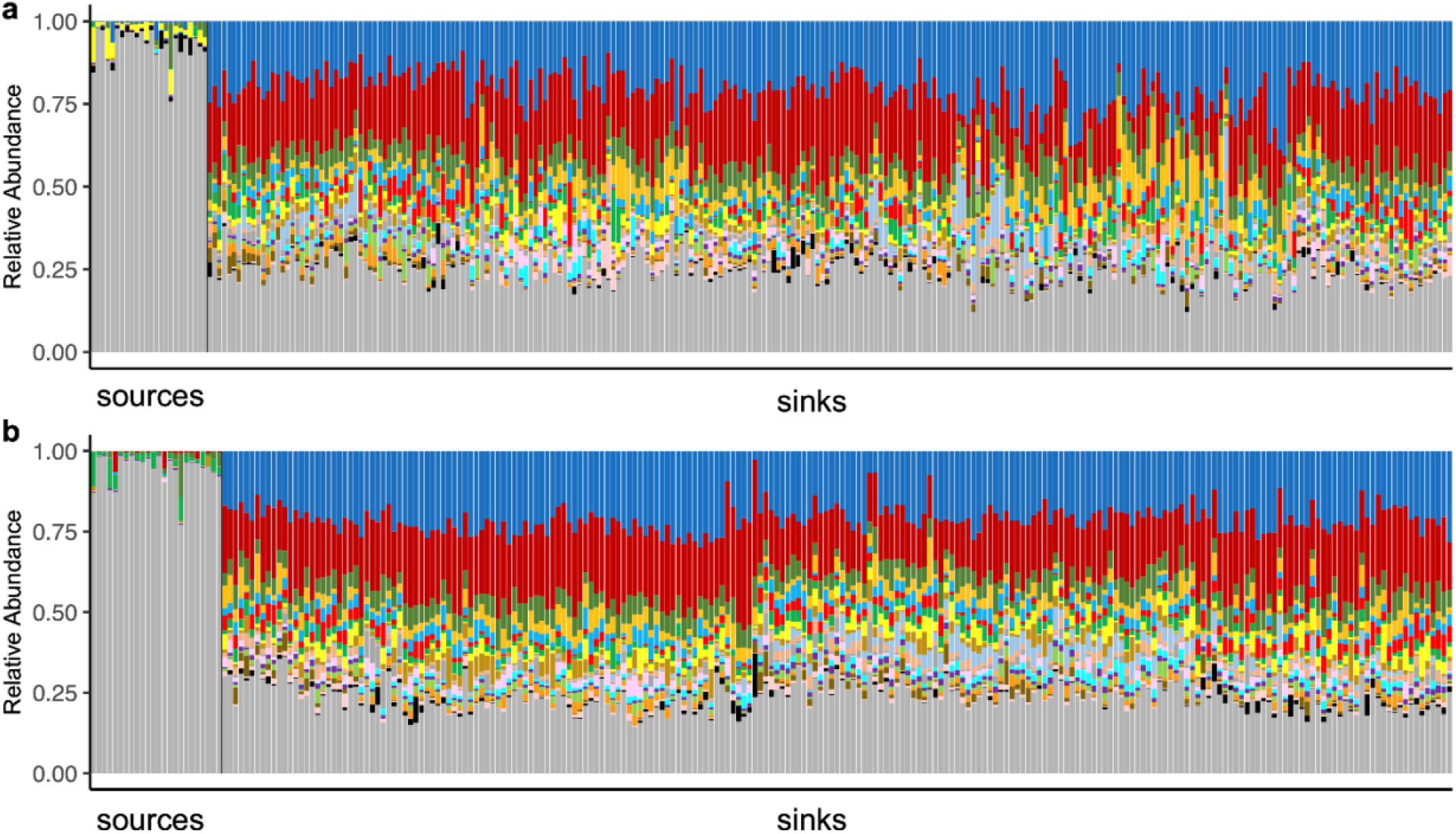
Taxonomic profiles of sources and sinks in community coalescence experiments. **a**, Pairwise mixing. **b**, Quadruple-wise mixing. For visualization purposes, we only showed the top-20 abundant ASVs. All other ASVs were grouped together and shown in gray. We found that some highly abundant ASVs in the source communities have very low abundances in the sink communities, whereas some low-abundance ASVs in source communities flourish in the sink communities. Also, a few ASVs in the sinks were not detected in the sources, indicating that either their relative abundances were below the detection limit or there was potential contamination.

**Figure S7:**
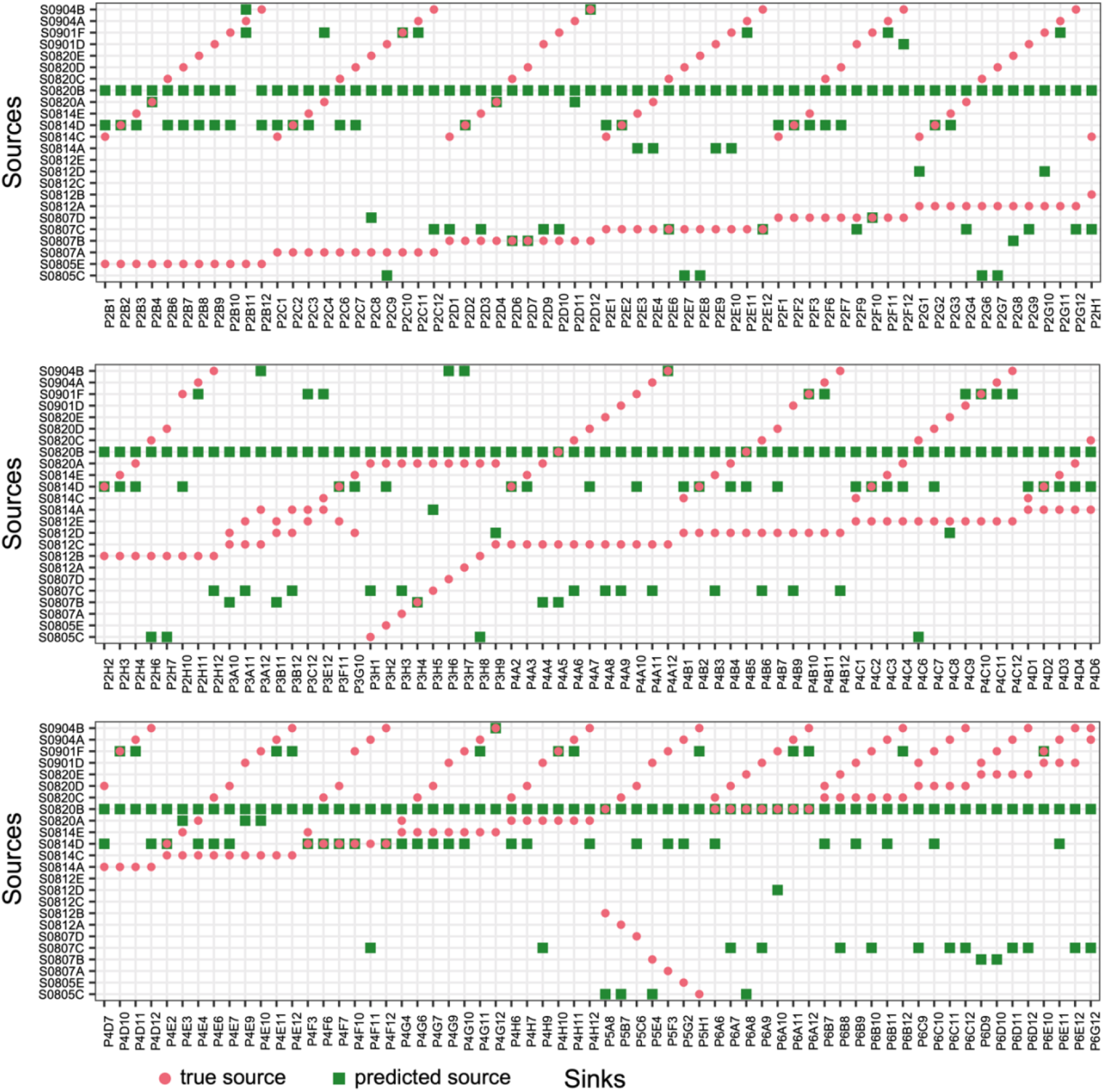
Performance of FEAST in identifying sources in pairwise community coalescence experiments. True sources (red cycles) vs. predicted sources (green squares) of each sink. For each sink, among the 24 known sources, the two sources with the top-two largest contributions predicted by FEAST were referred to as the predicted sources. In Fig.5, we showed the results of the first 64 sinks. Here we showed the results of the remaining 192 sinks.

**Figure S8:**
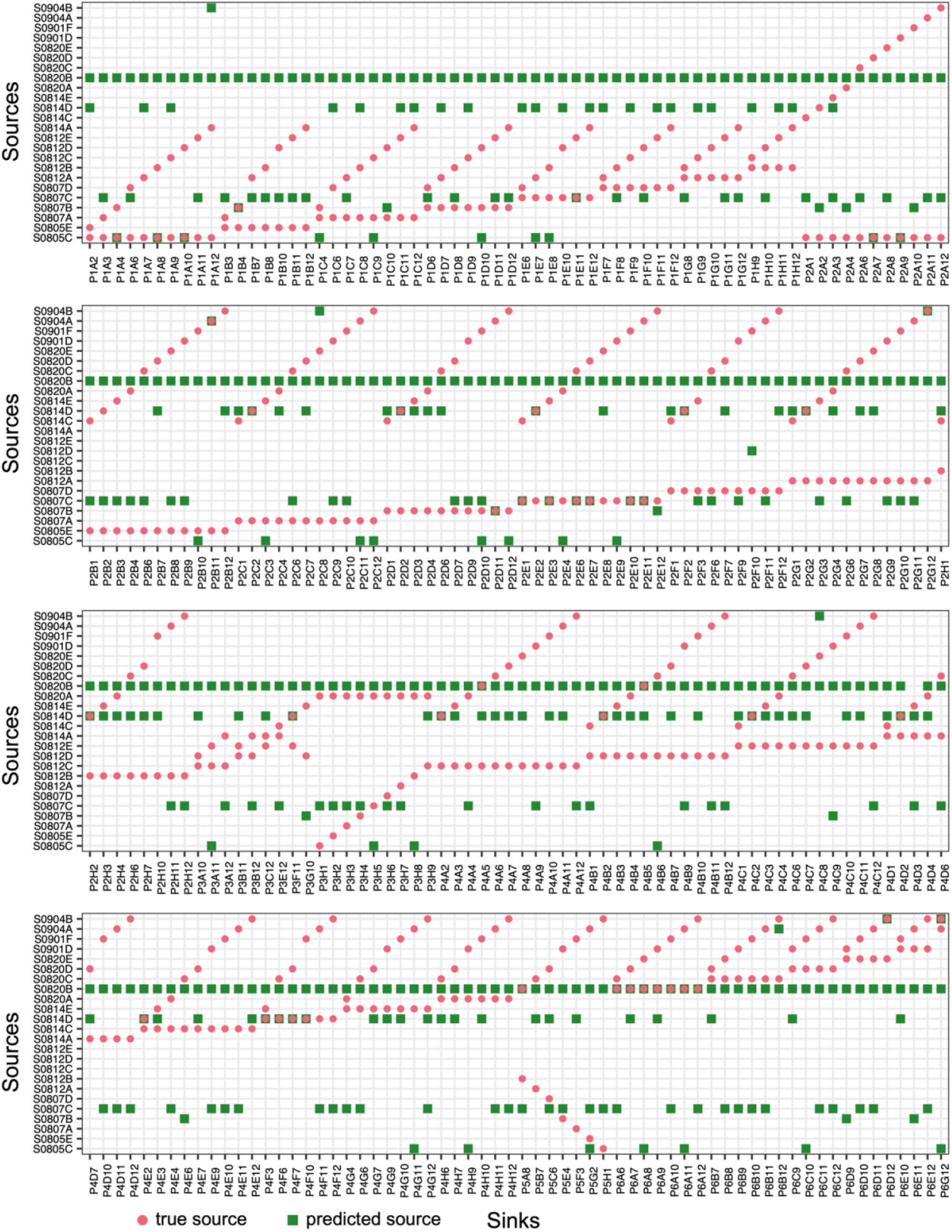
Performance of SourceTracker in identifying sources in pairwise community coalescence experiments. True sources (red cycles) vs. predicted sources (green squares) of each sink. For each sink, among the 24 known sources, the two sources with the top-two largest contributions predicted by SourceTracker were referred to as the predicted sources.

**Figure S9:**
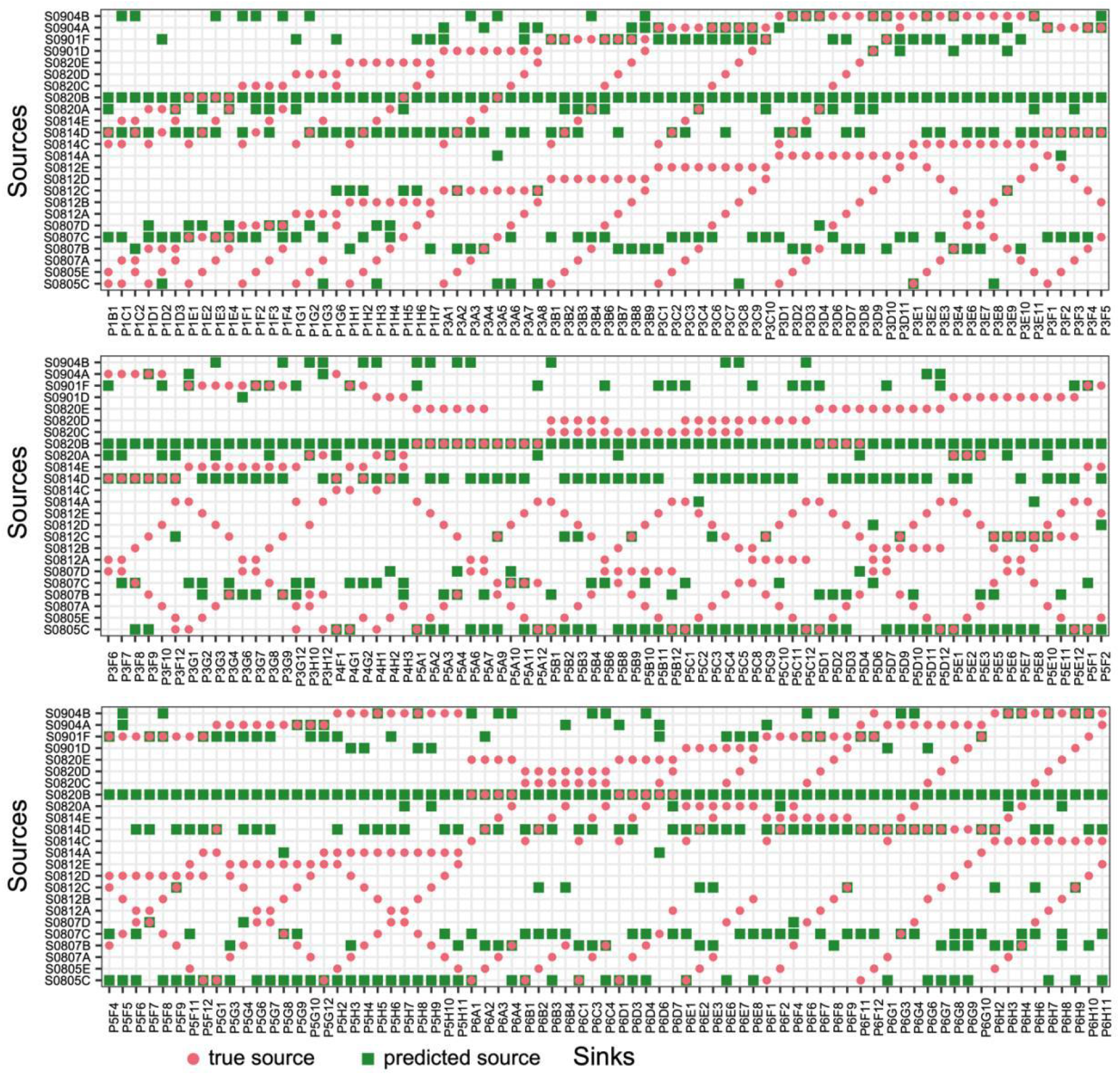
Performance of FEAST in identifying sources in quadruple-wise community coalescence experiments. There are 24 source communities (stool samples from 24 healthy individuals). Each sink community is obtained by mixing four different source communities *ex vivo*. The final composition of each sink was obtained from metagenomic sequencing of samples collected after 11 days of the *ex vivo* mixing. True sources (red cycles) vs. predicted sources (green squares) of each sink. (Each row includes 75 sinks). For each sink, among the 24 known sources, the four sources with the top-four largest contributions predicted by FEAST were referred to as the predicted sources.

**Figure S10:**
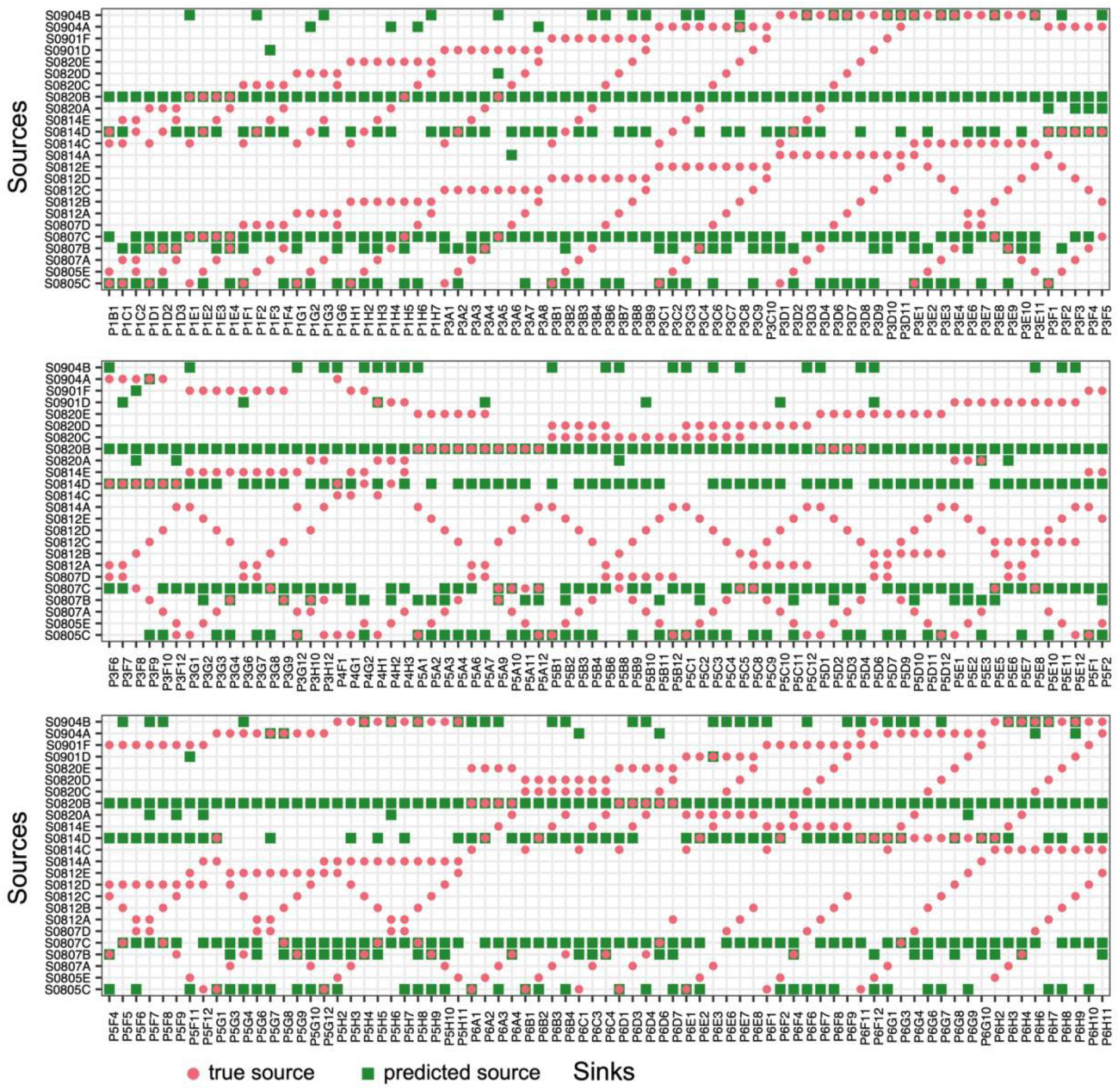
Performance of SourceTracker in identifying sources in quadruple-wise community coalescence experiments. There are 24 sources communities (stool samples from 24 healthy individuals). Each sink community is obtained by mixing four different source communities *ex vivo*. The final composition of each sink was obtained from metagenomics sequencing of samples collected after 11 days of the *ex vivo* mixing. True sources (red cycles) vs. predicted sources (green squares) of each sink. (Each row includes 75 sinks). For each sink, among the 24 known sources, the four sources with the top-four largest contributions predicted by SourceTracker were referred to as the predicted sources.

**Figure S11:**
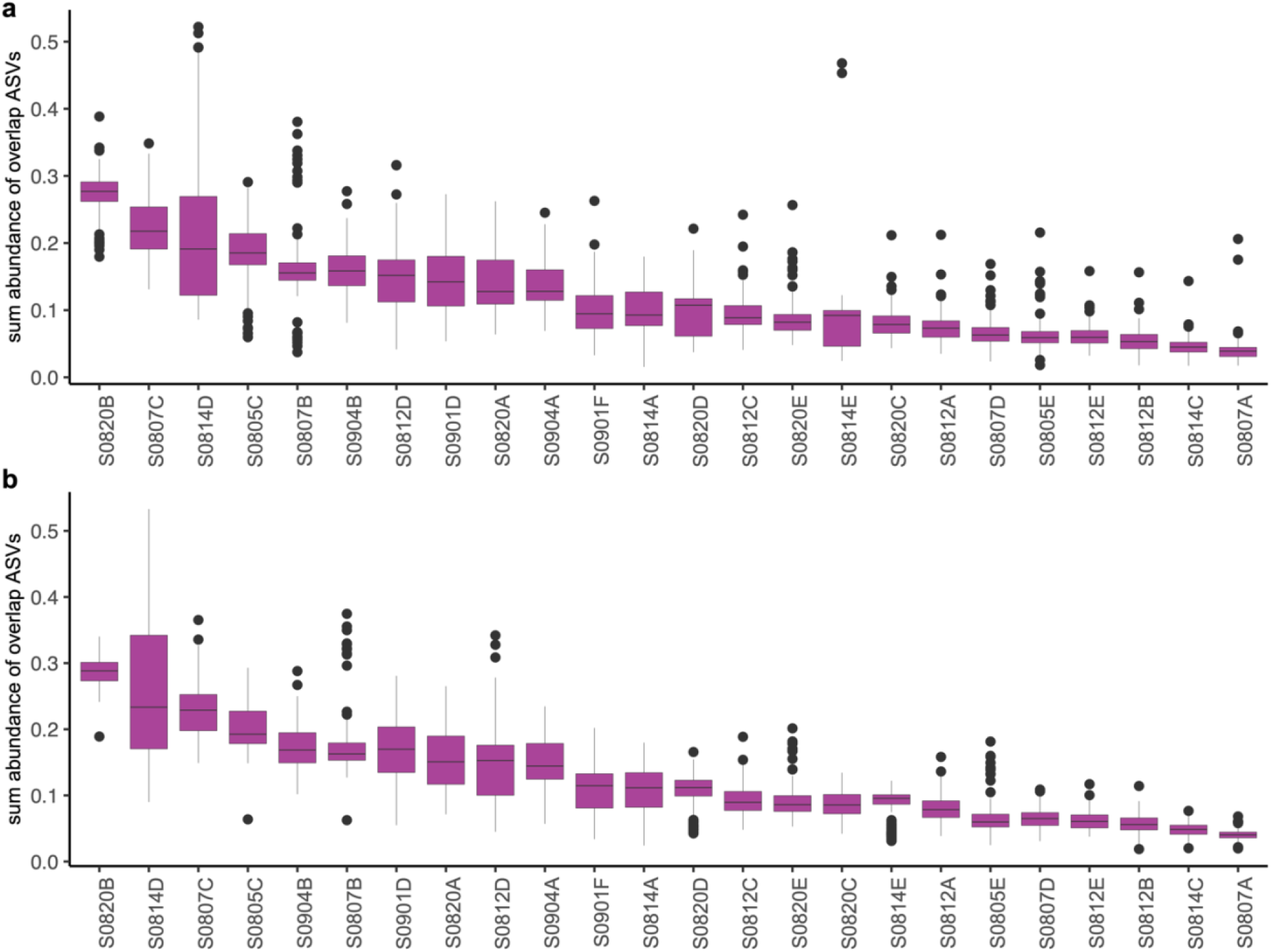
Relative abundances of common ASVs shared by sources and sinks. For each sink and source pair, we identified their common ASVs and calculated the total relative abundance of those common ASVs. Each boxplot represents the total relative abundance of common ASVs shared by this source and each of the 256 sinks in the pairwise community coalescence experiments (**a**); and 225 sinks in quadruple-wise community coalescence experiments (**b**).

